# Structural basis for Rab8a GTPase recruitment of RILPL2 *via* LRRK2 phosphorylation of switch 2

**DOI:** 10.1101/739813

**Authors:** Dieter Waschbüsch, Elena Purlyte, Prosenjit Pal, Emma McGrath, Dario R. Alessi, Amir R. Khan

## Abstract

Rab8a GTPase is associated with the dynamic regulation of membrane protrusions in polarized cells. Rab8a is one of several Rab-family GTPases that are substrates of leucine-rich repeat kinase 2 (LRRK2), a serine/threonine kinase that is linked to inherited Parkinson’s disease. Rab8a is phosphorylated at T72 (pT72) in its switch 2 helix and the post-translational modification facilitates phospho-Rab8a (pRab8a) interactions with RILPL2, which subsequently regulates ciliogenesis. Here we report the crystal structure of pRab8a in complex with the phospho-Rab binding domain of RILPL2. The complex is a heterotetramer with RILPL2 forming a central α-helical dimer that bridges two pRab8a molecules. The N-termini of the α-helices cross over to form an X-shaped cap (X-cap) that enables electrostatic interactions between Arg residues from RILPL2 and the phosphate moiety from pT72. RILPL2 residues in the X-cap that are critical for pRab8a binding are conserved in the RILP family of effector proteins. We find that JIP3 and JIP4 also interact specifically with LRRK2-phosphorylated Rab10, suggesting a general mode of recognition for phosphorylated Rab GTPases by phospho-specific effectors.

## Introduction

Parkinson’s disease (PD) is a disorder of the central nervous system that manifests as a progressive degeneration of motor mobility, loss of balance, and tremors. Features of the pathology include loss of dopaminergic neurons in the midbrain and the presence of protein aggregates termed Lewy bodies, composed mainly of α-synuclein, in surviving neurons [1]. About 10% of cases have a genetic basis, with the most common gene being the *Leucine-Rich Repeat Kinase 2* (*LRRK2)*[2],. The gene product is a 2,527-residue (286kDa) protein with multiple domains belonging to the ROCO family that is involved in regulation of autophagy, mitochondria, and Golgi dynamics [3]. The kinase domain, located near the C-terminus, phosphorylates itself and other proteins at serine/threonine residues [2, 4, 5]. Preceding the kinase domain, there is a Ras-like ROC domain (<u>R</u>as <u>o</u>f <u>c</u>omplex) followed in tandem by a COR domain (<u>C</u>-terminal <u>o</u>f <u>R</u>as). The ROC domain binds nucleotides (GTP/GDP) and is distantly related to the Rab family of small GTPases. The ROC-COR tandem domains regulate LRRK2 activity and numerous missense mutations have been localized to these regulatory and kinase domains [6–8]. In addition to early onset forms of PD associated with autosomal dominant mutations, LRRK2 is also linked to late-onset sporadic cases of PD [9].

Insight into LRRK2 functions has progressed significantly with the finding that a subset of small GTPases that include Rab8 and Rab10 are physiological substrates of the enzyme [5, 10]. Rabs comprise the largest group (∼70 members) of the Ras superfamily, and they cycle between an active GTP-bound and inactive GDP form to regulate membrane trafficking in eukaryotic cells [11]. The nucleotide-bound state of Rabs is regulated by GTPase activating proteins (GAPs) and GTP/GDP exchange factors (GEFs), and active Rabs migrate to distinct sub-cellular compartments where they recruit cytosolic effector proteins. The ‘switch’ regions of Rabs, termed switch 1 and 2, undergo local conformational changes that enable recruitment of GTP-specific effectors, which subsequently control processes such as vesicle formation/fusion, motility, and other aspects of cell dynamics [12]. LRRK2 phosphorylates Rab8a at T72 and Rab10 at T73, conserved threonine residues located on the α-helical switch 2 region. This post-translational modification modulates interactions between Rabs and their binding partners [5, 13]. For example, it inhibits interaction with some effectors such as Rab GDP dissociation inhibitors (GDIs) and Rabin-8, a GEF for Rab8a [5]. LRRK2 phosphorylation of Rab8a and Rab10 also promoted interaction with two poorly studied scaffolding proteins termed RILPL1 (Rab interacting lysosomal like protein 1) and RILPL2 [13], that were previously implicated in regulating ciliogenesis[14]. RILPL1 and RILPL2 belong to the RILP (Rab interacting lysosomal protein) family of effector proteins[15]. Unlike RILPL1 and RILPL2, RILP was not observed to interact with LRRK2 phosphorylated Rab8a or Rab10[13], however, it is a known effector for Rab7a GTPase[16]. A recent study has suggested that RILP may bind more strongly to Rab7a phosphorylated at the equivalent site to LRRK2 by the related LRRK1 kinase[17]. Cellular studies have confirmed that LRRK2 blocks ciliogenesis by phosphorylating Rab8a and Rab10 and promoting RILPL1 interaction[18]. RILPL1 and RILPL2 are homologous and were shown to interact with LRRK2 phosphorylated Rab8a and Rab10 via a C-terminal phospho-Rab binding domain that encompasses a conserved region of the protein. This region of the protein is also known as the RILP homology domain 2 (RH2) and encompasses residues 291-356 on RILPL1 and residues 130-201 on RILPL2. RILPL1 and RILPL2 also contain an N-terminal RH1 domain that binds to the globular tail domain (GTD) of myosin Va[19].

Upstream of LRRK2 it has been shown that Rab29 recruits LRRK2 to the Golgi and activates the kinase, leading to an enhanced phosphorylation of downstream Rab GTPases, as well as increased autophosphorylation [20–22]. Rab32 is not a target for the kinase but it interacts with LRRK2 and regulates its sub-cellular localization [23]. Thus, LRRK2 is at the center of a Rab signaling cascade that is a key to understanding the biological functions of LRRK2 and its relationship to PD. Here we describe the crystal structure of T72 phosphorylated GTP bound form of Rab8a in complex with a minimal phospho-Rab binding domain of RILPL2 at 1.8 Å resolution. The structure reveals that the phosphothreonine (pT72) is recognized by a conserved arginine from the RILP family of proteins, suggesting a general mechanism for phospho-specific recognition of effectors by Rab GTPases.

## Results and Discussion

### Overall structure of pRab8a:RILPL2 complex

For these studies we utilized a mutant of the globular G-domain of Rab8a (Q67L, residues 1-181) that binds GTP constitutively. Full-length RILPL2 complexes failed to crystallize, but the phospho-Rab binding domain of RILPL2 (residues 129 to 165) yielded crystals in complex with pRab8a. This region has an N-terminal hexahistidine tag and is the minimal fragment with high sequence similarities to all members of the RILP effector family. The complex of pRab8a(GTP):RILPL2 is organized as a heterotetramer in the asymmetric unit (Fig 1A and 1B), with a central parallel α-helical dimer of the phospho-Rab binding domain of RILPL2 bridging two molecules of pRab8a *via* hydrophobic and polar interactions. Both molecules of pRab8a in the complex have GTP in the nucleotide pocket and their switch 1 and 2 conformations resemble the structure of active Rab8a [PDB code 4lhw [24]]. Each pRab8a molecule interacts with both α-helices of the effector, burying approximately 625 Å^2^ of surface area at each interface. The dual α-helical interactions are restricted to the N-terminal segment of RILPL2. As the coiled coil extends toward the C-termini, a single α-helix interacts with each Rab monomer by interfacing with switch 1 and strand β2 of the interswitch region (Fig 1C). The topology of the complex on Golgi membranes would be consistent with the N-terminus of RILPL2 (1-128) oriented above the complex in the orientation shown in Fig 1A to enable interactions with the GTD of myosin Va. The C-termini of pRab8a (177-207) and RILPL2 (160-211) would reside proximal to the membrane, as indicated with dashed lines following helix α5 of pRab8a. In the ensuing discussions, the acronyms ‘RL2’ and ‘R8’ will be used in superscript format to denote RILPL2 and Rab8a residues, respectively.

**Figure 1:**
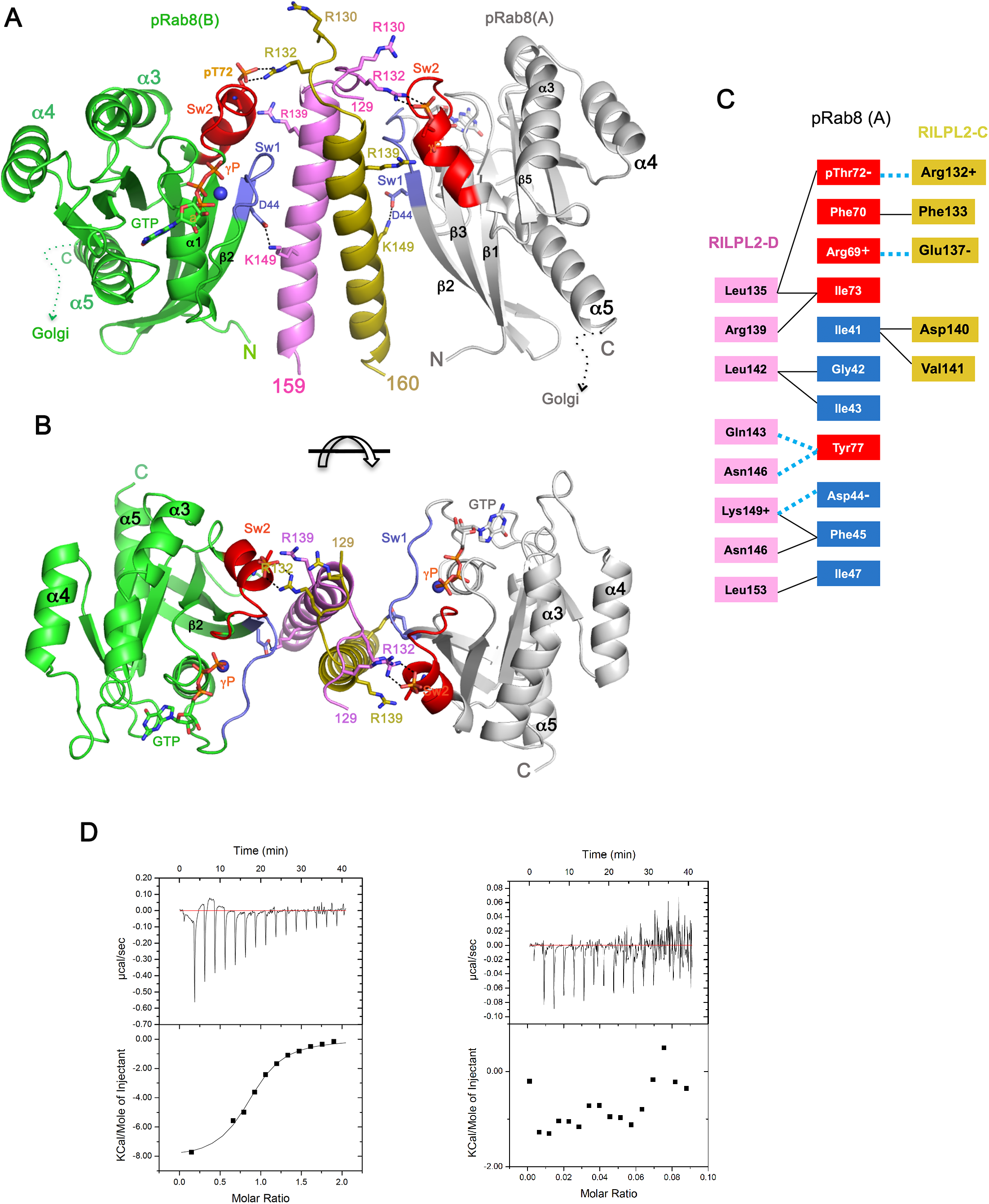
Structure of pRab8a in complex with phospho-Rab binding domain of RILPL2. **A**, heterotetrameric assembly of two pRab8a molecules bridged by a central α-helical dimer of the phospho-Rab binding domain of RILPL2 (129-165). The two chains of RILPL2 are in magenta and dark yellow. For pRab8a, switch 1 is shown in blue, switch 2 in red. **B**, view of the complex down the two-fold axis of the heterotetramer, 90° relative to orientation in A. **C**, cartoon representation of the contacts between one molecule of Rab8a and the dimer of RILPL2. Polar interactions are indicated in dotted blue lines. **D**, isothermal titration calorimetry analyses of the interactions between pRab8a and phospho-Rab binding domain of RILPL2. *Left*, titration of RILPL2 (residues 129-165) into pRab8a(GTP). *Right*, titration of RILPL2 into Rab8a(GTP).

The affinity of the interaction between pRab8a and the phospho-Rab binding domain (129-165) of RILPL2 was evaluated by isothermal titration calorimetry (Fig 1D, *left*). The experiment revealed a K_d_ = 3.3μM (±0.5) for the interaction, which indicates a relatively weak affinity that is similar to other physiological Rab:effector complexes [12]. There were no detectable interactions between non-phosphorylated Rab8a(GTP) and RILPL2 (Fig 1D, *right*).

### Phosphorylated switch 2 of Rab8a interacts with an ‘X-cap’ region of RILPL2

The N-termini of the RILPL2 dimer (N129^RL2^-T134^RL2^) cross over in an extended conformation preceding the first α-helical turn, forming an X-shaped cap (X-cap) over the coiled coil (Figs 2A and 2B). This X-cap contains two conserved arginines (R130^RL2^, R132^RL2^) that previous mutagenesis analysis showed were required for interaction with pRab8a [13]. Intimate contacts within the X-cap include reciprocal backbone hydrogen bonds between residues R132^RL2^-Phe133^RL2^ that resemble a short anti-parallel β-sheet (Fig 2B). The phosphate moiety from pT72 interacts with the guanidino group (NH1/NH2) of R132^RL2^ on both sides of the symmetric complex, with O/N distances between 2.5-2.9 Å (Fig 2C). Thus, R132^RL2^ binds directly to the phosphorylated threonine, and this residue is conserved in the RILP family of effectors (Fig 4C). In contrast to these direct electrostatic and hydrogen-bonding interactions, the side chain of R130^RL2^ is relatively disordered and more distant from pT72 (>6 Å). An electrostatic surface map of RILPL2 reveals the strongly positive charges at the X-cap that enable recognition of pRab8a (Fig 2D). In addition to electrostatic contacts, the X-cap residue F133^RL2^ contributes to a complementary hydrophobic interface with F70^R8^ and I73^R8^ from switch 2 (Fig 2B). The side chain of T134^RL2^ acts as a capping residue by nucleating the α-helix *via* a hydrogen bond (3.2 Å) to the backbone NH of E137^RL2^. Therefore, the term X-cap is appropriate for this region of RILPL2.

**Figure 2:**
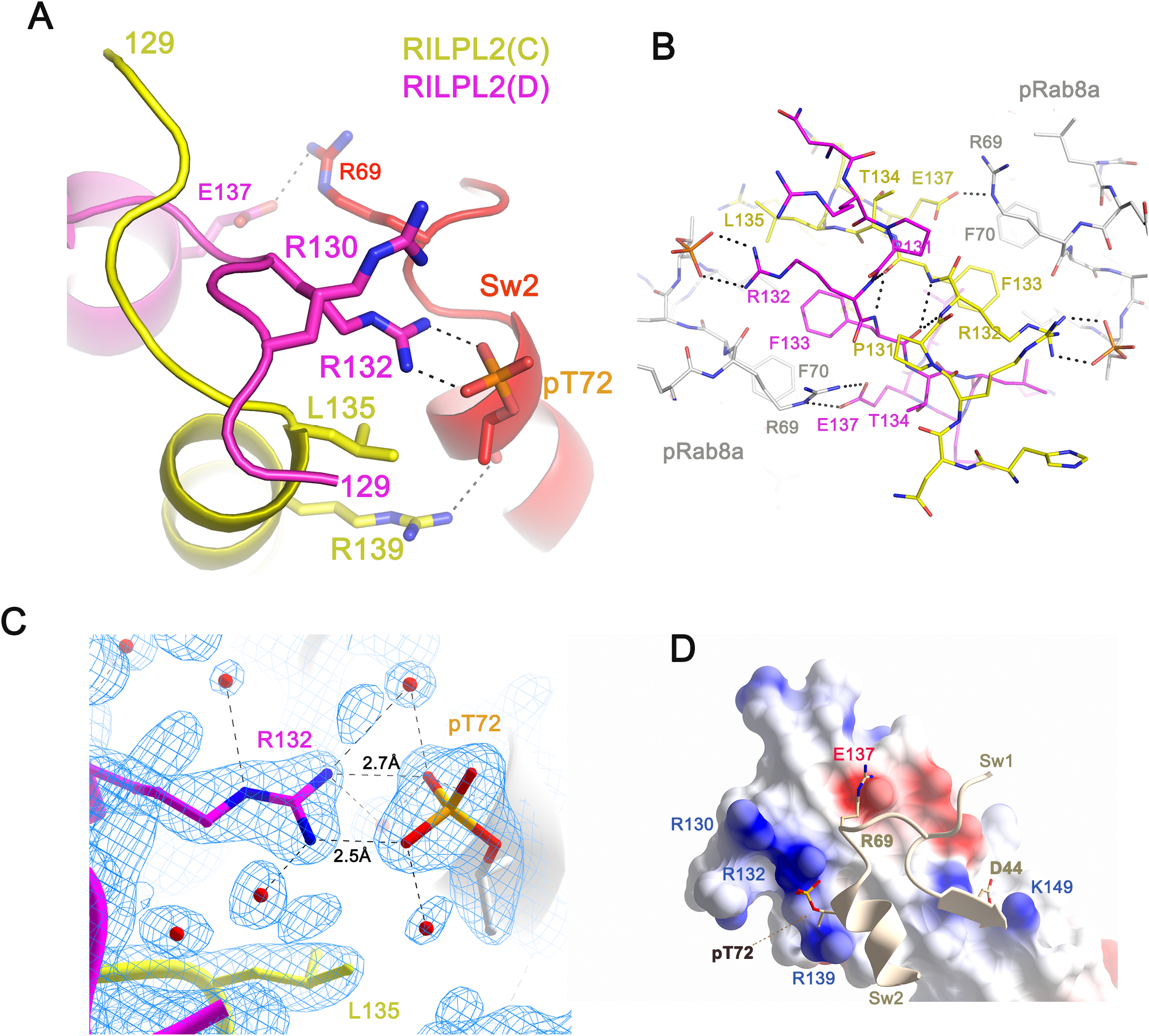
Structural details of pT72 recognition by the X-cap of RILPL2. **A**, view of an interface between Rab8a and the dimer of RILPL2. **B**, stick model of the interactions at the X-cap of RILPL2. **C**, electron density (2Fo-Fc, 1.2σ) at the site of pT72 (chain B) binding to R132^RL2^ (chain D). The side chain of L135^RL2^ from chain C of RILPL2 lies within van der Waals contact (4 Å) of the β-branched methyl group of pT72. **D**, electrostatic surface rendering of the X-cap. Blue is positive and red is negative, while switch 1 and 2 of pRab8a are ribbons with key residues represented as sticks.

### Mutational analyses of the binding interface

The contribution of RILPL2 residues to complex formation with pRab8a was evaluated by mutagenesis. LRRK2-phosphorylated Rab8a was subjected to co-immunoprecipitation studies in cells using wild type and mutant forms of full length RILPL2. For these studies a pathogenic LRRK2[R1441G] mutant was overexpressed to ensure maximal phosphorylation of Rab8a. We also treated cells with and without a potent and selective LRRK2 inhibitor termed MLi-2 [25] for 90 min to induce dephosphorylation of Rab8a and, therefore, block association with RILPL2 (Figs 3 and EV3A-B). These results revealed that all mutations of R132^RL2^, which directly interacts with pT72, abolished the interaction with pRab8a in cells. The exquisite specificity of this contact is reflected by the R132K^RL2^ mutation, which was sufficient to abolish the interaction (Fig 3). Mutation of K149^RL2^ - which forms a salt bridge with D44^R8^ (switch 1) - also abolishes the interaction between RILPL2 and pRab8a. This interaction, as well as hydrophobic packing of I41^R8^ against the RILPL2 α-helical dimer, presumably encodes GTP-dependent switch 1 specificity. Modest effects were observed with mutations of R139^RL2^ and L135^RL2^, which form contacts with switch 2 adjacent to the R132^RL2^:pT72 interaction. L135 packs against the β-branched methyl substituent of pT72, while R139 forms a hydrogen bond with the backbone carbonyl oxygen of pT72 (Figs 2A and 2C).

**Figure 3:**
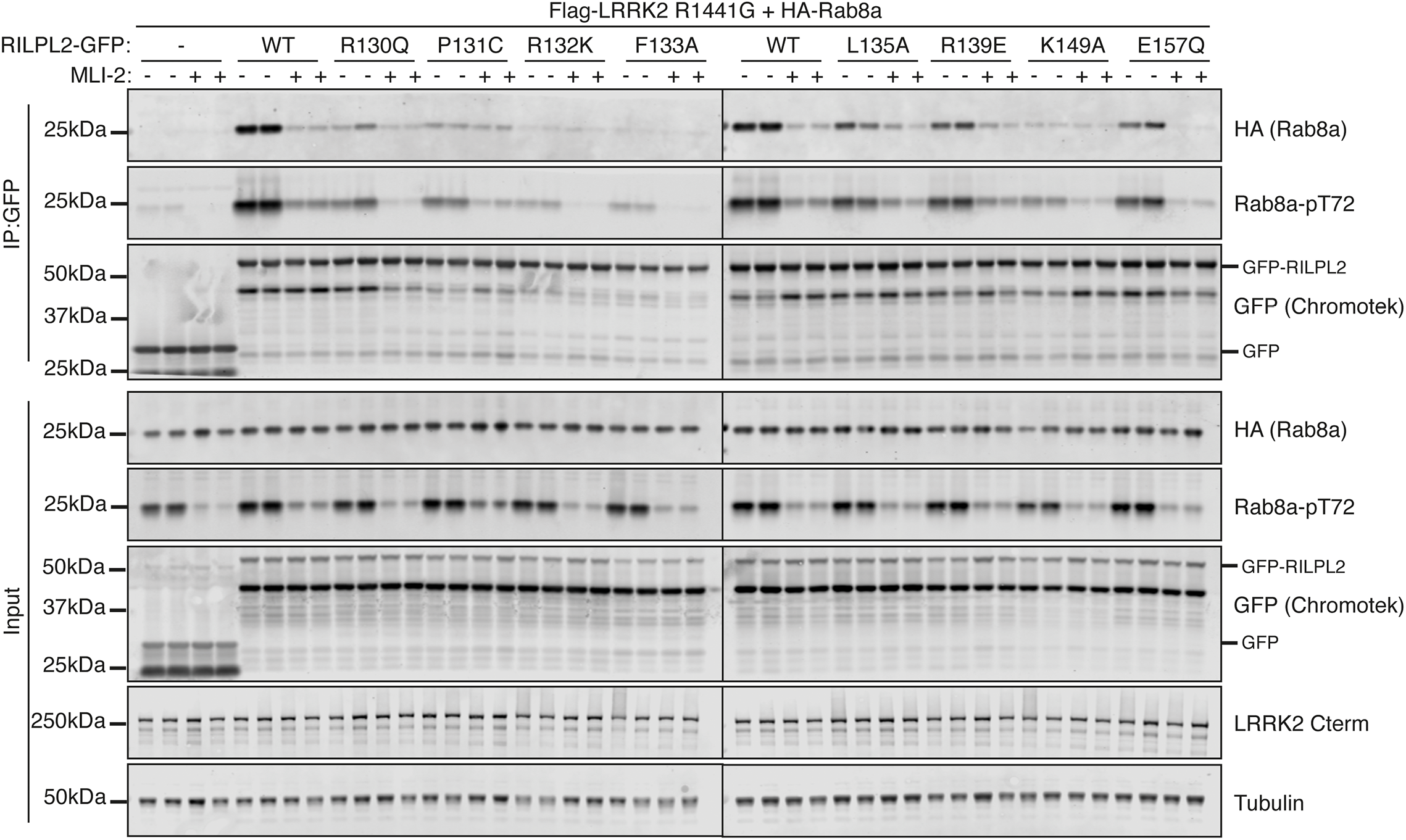
Mutational analyses reveal hotspots of pRab8a:RILPL2 interactions. HEK293 cells were transiently transfected with constructs expressing the indicated components. 48 h post-transfection, cells were treated with ± 500 nM MLi-2 for 90 min and then lysed. Upper-panel labelled IP:GFP, RILPL2-GFP was immunoprecipitated using GFP binder sepharose and immunoprecipitates evaluated by immunoblotting with the indicated antibodies. Immunoblots were developed using the LI-COR Odyssey CLx Western Blot imaging system analysis with the indicated antibodies at 0.5-1 µg/mL concentration. Lower-panel labelled Input-10 µg whole cell lysate was subjected to LI-COR immunoblot analysis. Each lane represents cell extract obtained from a different dish of cells. Similar results were obtained in two separate experiments.

**Figure 4:**
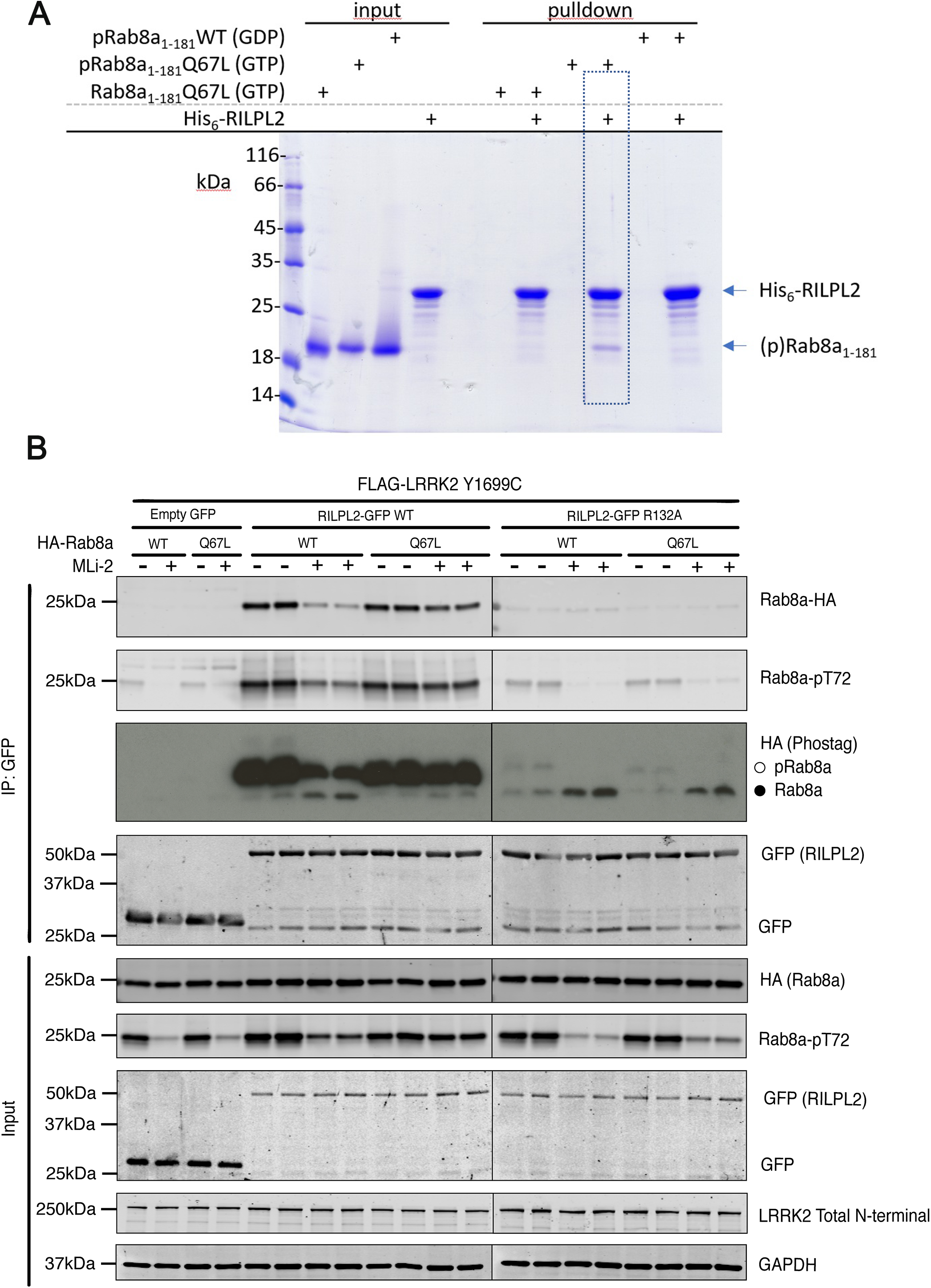
Evidence that RILPL2 binds to the GTP bound conformation of phosphorylated Rab8a in cells. **A**, Direct *in vitro* pulldowns were performed using purified His_6_-tagged RILPL2 (*full length*) as bait and untagged Rab8a as prey. Rab8a species were either non-phosphorylated Rab8a-GTP(Q67L), pRab8a-GTP(Q67L), or pRab8a(GDP). Protein concentrations were 10µM for bait and prey, inputs are 2 µg; n≥3, Coomassie stain for visualization. Dotted lines emphasize that only pRab8a(GTP) binds to RILPL2. **B**, HEK293 cells were transiently transfected with constructs expressing the indicated components. 24 h post-transfection, cells were treated with ± 100 nM MLi-2 for 90 min and then lysed. Upper-panel labelled IP:GFP, RILPL2-GFP was immunoprecipitated using GFP binder sepharose and immunoprecipitates evaluated by immunoblotting with the indicated antibodies. Immunoblots were developed using the LI-COR Odyssey CLx Western Blot imaging system analysis with the indicated antibodies at 0.5-1 µg/mL concentration. Lower-panel labelled Input-10 µg whole cell lysate was subjected to LI-COR immunoblot analysis. Each lane represents cell extract obtained from a different dish of cells. Similar results were obtained in two separate experiments.

One apparent inconsistency between the structure and the mutagenesis experiments is the contribution of R130^RL2^ toward complex formation. Mutational studies reveal that R130Q^RL2^ (Fig 3), R130E^RL2^ (Fig EV3A) and R130A^RL2^(Fig EV3A) severely compromise the interactions with pRab8a. As previously mentioned, the side chain of R130^RL2^ is relatively disordered and lies more than 6 Å away from pT72. However, modeling of alternate conformations of the side chains brings the guanidino group to within 5 Å of the phosphate on both interfaces. Thus, a reasonable explanation is that Arg130^RL2^ contributes to pT72 through long-range electrostatic interactions with the phosphate.

### GTP dependency of the interaction between pRab8a and RILPL2

The GTP dependency of pRab8a interactions with full-length RILPL2 were investigated using *in vitro* pulldowns (Fig 4A). The interaction with RILPL2 *in vitro* is dependent on both the GTP conformation and phosphorylated T72 for Rab8a. Non-phosphorylated Rab8a and pRab8a(GDP) did not interact measurably with RILPL2.

To further investigate the GTP dependency in cells, we co-expressed RILPL2 with either wild type Rab8a, Rab8a[Q67L] (GTP trapped conformation) or Rab8a[T22N] (GDP bound conformation) in the presence of pathogenic LRRK2[Y1699C] to induce maximal Rab8a phosphorylation (Fig 4B and EVFig4). For these experiments, cells were treated ± LRRK2 MLi-2 inhibitor and phosphorylation of wild type and mutant Rab8a was assessed in cell extracts as well as RILPL2 immunoprecipitates. Strikingly, this revealed that both cell lysates and immunoprecipitates of the pRab8a[Q67L] GTP locked mutant, MLi-2 failed to induce dephosphorylation of Rab8a over a 90 min period (Fig 4). We interpret this result to indicate that the pRab8a[Q67L] GTP locked conformation remains stably associated with RILPL2 over this period and is thus protected from dephosphorylation by cellular phosphatases. In comparison, MLi-2 treatment induced significant dephosphorylation of wild type Rab8a that is presumably interconverting between the GTP and GDP conformation (Fig 4). The GDP-locked conformation of Rab8a[T22N] was expressed at much lower levels in cell than wild type or Rab8a[Q67L] (EVFig4). Nevertheless, no association of RILPL2 with, Rab8a[T22N] was observed. Furthermore, phosphorylation of Rab8a[T22N] was not observed which is consistent with the GTP bound conformation being regulated by LRRK2.

### X-cap binding to phosphorylated switch 2 may be a general mode of recognition

The structure of the Rab binding domain of RILP bound to Rab7 has been determined previously [Fig 5A; [26]]. Similar to RILPL2, the effector forms a parallel coiled coil that binds two Rab7 molecules as a heterotetrameric complex. RILP does not have an ‘X-cap’ at the N-terminus – the first few residues in the structure beginning at C241 are disordered until the start of the core helix at Glu246 (Figs 5A-B). Following helix α1 of the Rab binding domain of RILP, a loop brings a second anti-parallel helix α2 back toward the α1 dimer and is stabilized by hydrophobic interactions. Helix α2 is unlikely to be conserved in RILPL1/2 given a string of glycine and proline residues in these proteins following helix α1, which would be disruptive to secondary structures (Fig 5C).

**Figure 5:**
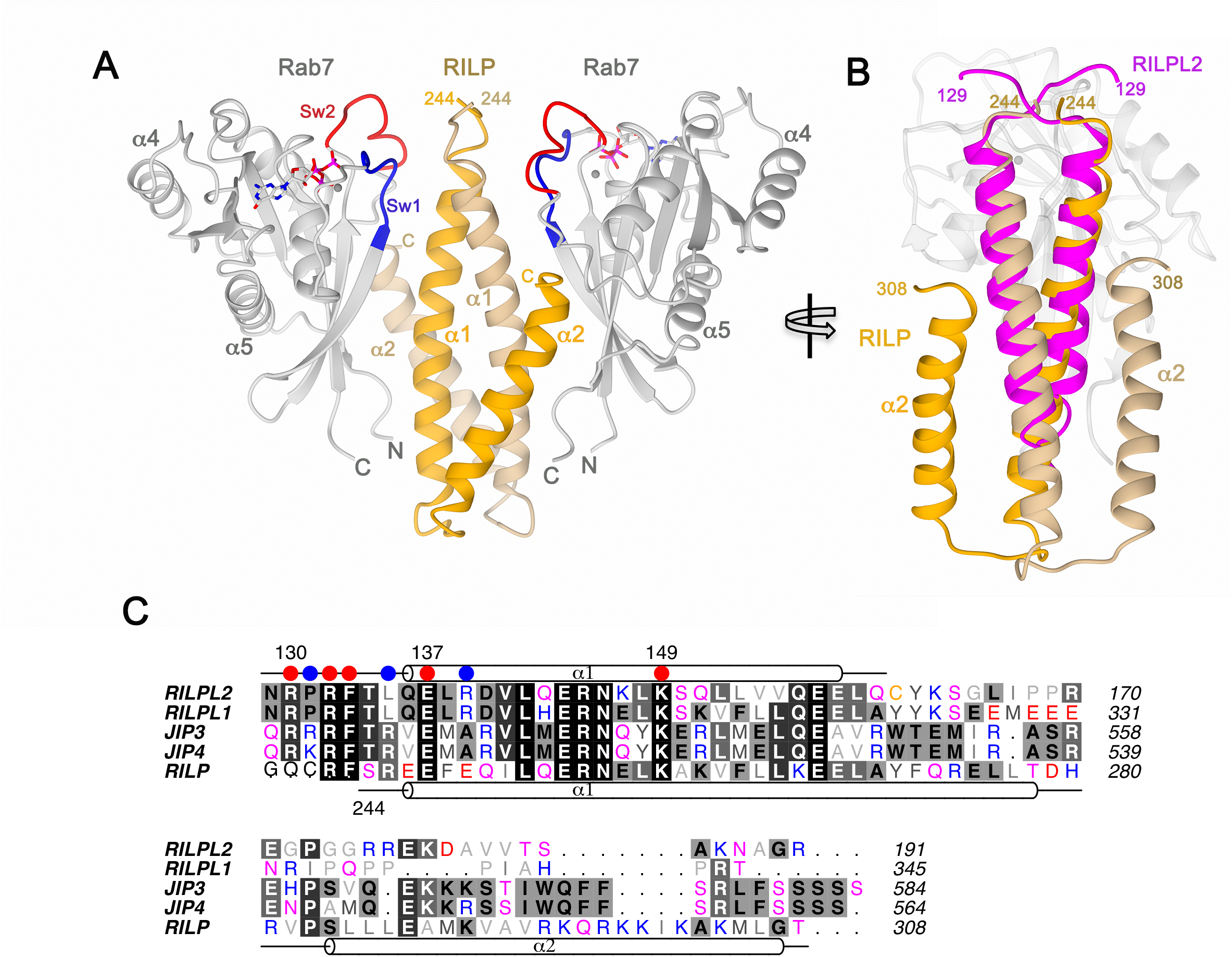
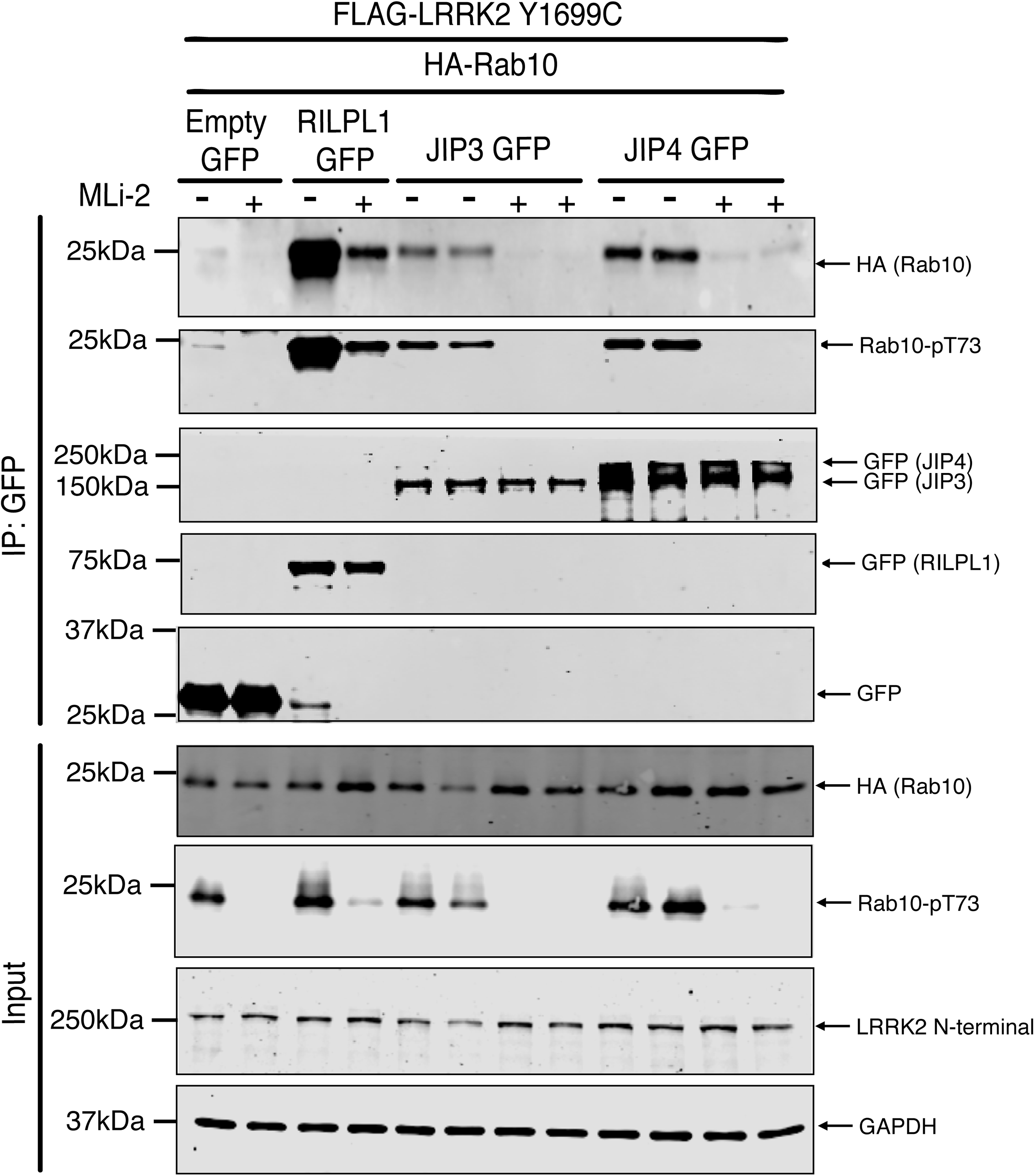
Sequence and structural comparisons of Rab complexes with the RILP-like family of effectors. **A**, Structure of Rab7 in complex with the Rab-binding domain of RILP. **B**, Superposition of RILPL2 onto a single binding interface of Rab7:RILP, showing conservation of the α-helical coiled coil. The figure is rotated 90° along the horizontal axis, relative to A. **C**, sequence alignment of the RILP-like family of proteins. Positively charged residues in the phospho-Rab binding domain of RILPL2 that are exquisitely sensitive to mutagenesis for pRab8a:RILPL2 complex formation have red circles above them. R139 (orange circle) has a partial defect in binding following mutation to glutamate. **D**, Evidence that JIP3 and JIP4 bind to LRRK2 phosphorylated Rab10 in cells. HEK293 cells were transiently transfected with constructs expressing the indicated components. 24 h post-transfection, cells were treated with ± 100 nM MLi-2 for 90 min and then lysed. Upper-panel labelled IP:GFP, RILPL2-GFP, JIP3-GFP, JIP4-GFP was immunoprecipitated using GFP binder sepharose and immunoprecipitates evaluated by immunoblotting with the indicated antibodies. Immunoblots were developed using the LI-COR Odyssey CLx Western Blot imaging system analysis with the indicated antibodies at 0.5-1 µg/mL concentration. Lower-panel labelled Input-10 µg whole cell lysate was subjected to LI-COR immunoblot analysis. Each lane represents cell extract obtained from a different dish of cells. Similar results were obtained in two separate experiments.

The hotspots for pRab8a:RILPL2 interactions, as gleaned from cellular interactions with RILPL2 mutants, are shown with circles above the sequences of RILP family proteins (Fig 5C). The red circles denote essential residues for recognition of pRab8a(GTP) and are mostly conserved in the phospho-Rab binding domains. R139^RL2^ mutations have modest effects on Rab8a recognition (Figs 3 and EVFig3), and are not conserved (blue circle). P131^RL2^ is variable in this family and the mutant P131A^RL2^ does not affect binding, suggesting that sequence variability is possible within the X-cap. Interestingly, LRRK1 phosphorylation of Rab7 at Ser72 (switch 2) has recently been shown to promote interactions with RILP[17]. The RILP construct used for recombinant expression begins at C241 and the PDB file (1yhn) for the complex RILP:Rab7 contains the co-ordinates for RILP beginning at S244 (T134^RL2^). It is possible that an X-cap could form and bind to pS72 (Rab7) given a modest extension of the RILP polypeptide toward the N-terminus to stabilize backbone anti-parallel hydrogen bonds. Although RILP lacks a positive residue (Q240), the structure of pRab8a:RILPL2 reveals that Arg130^RL2^ is distant from pT72 and may be tolerated as a glutamine within a possible X-cap in the RILP:pRab7 complex.

Sequence comparisons suggest that two other scaffolding proteins previously implicated in JNK signaling, namely JIP3[27] and JIP4[28], likely possess a phospho-Rab binding domain (Fig 5C). These proteins have key residues equivalent to R130^RL2^, R132^RL2^ and K149^RL2^ in their putative phospho-Rab binding domain (Fig 5C). We therefore tested whether full length JIP3 or JIP4 would bind to LRRK2-phosphorylated Rab8a or Rab10 in cells employing the co-expression assay utilized above to assess interaction of pRab8a with RILPL2 (Fig 5D and EVFig5). These results revealed that both JIP3 and JIP4 specifically associated with LRRK2-phosphorylated Rab10. Addition of MLi-2 ablated interaction of JIP3 and JIP4 with Rab10 consistent with the interaction being phosphorylation dependent (Fig 5D). In contrast, a weak interaction just above background was observed between Rab8a and JIP3/4, which was not dependent upon phosphorylation by LRRK2 (Fig EVFig5). This emphasizes that phospho-Rab binding domains are likely to display selectivity for different phosphorylated Rab proteins. Further work is required to identify which sets of phospho-Rab proteins interact with JIP3/4 and how specificity for Rab GTPases is achieved.

In summary, the crystal structure of pRab8a in complex with RILPL2 provides molecular details of how post-translational modifications of Rab GTPases enable effector specificity. The structure reveals a positively charged X-cap at the N-termini of the RILPL2 α-helical dimer that mediates interactions with phosphothreonine in switch 2. The residue pT72 is situated within the switch/interswitch region and thus provides a means of tuning the strength of interactions with GAPs, GEFs and other effectors to regulate membrane trafficking upon LRRK2 activation in cells.

## Materials and Methods

### Protein expression and purification

The cDNA for Rab8a (residues 1-181, Q67L) lacking the flexible C-terminal tail was ordered from Genscript in a codon-optimized form to enable *E.coli* expression. The cDNA was cloned into pET28a at the NdeI/BamH1 sites for both of these constructs. The Rab8a *wildtype* (WT) construct was made by site directed mutagenesis using the following primers: 5’-GG GAT ACC GCG GGT <u>CAG</u> GAA CGT TTT CGT AC-3’ (for) and: 5’-GT ACG AAA ACG TTC <u>CTG</u> ACC CGC GGT ATC CC-3’ (rev). Expression was carried out in 2xYT Broth supplemented with 34 μg/ml kanamycin (FORMEDIUM™) at 37 °C. At an OD_600_ of 0.7 the culture was induced with 0.5 mM IPTG (FORMEDIUM™), after which cells were grown for a further 4 hours at 37 °C or 18 °C overnight. Cells were harvested by centrifugation and the pellets were resuspended in His-tag extraction buffer (20 mM Tris-HCl, 300 mM NaCl, 5 mM MgCl_2_, 20 mM imidazole and 10 mM β-mercaptoethanol, pH 8.0) along with 0.5 mM PMSF protease inhibitor (Sigma). Cells were lysed by sonication and the cell lysate was centrifuged at 26,000 x *g* for 45 minutes at 4 °C to remove cellular debris. The supernatants were filtered and loaded onto a nickel agarose resin (QIAGEN). The resin was washed with a 10-fold excess of extraction buffer and 5-fold excess wash buffer (extraction buffer supplemented with 40 mM Imidazole). The protein was eluted using extraction buffer supplemented with 200 mM imidazole. The cDNA corresponding to full-length RILPL2 (residues 1-211) was synthesized in codon-optimized format for *E.coli* expression and inserted into the Nde1/BamH1 site of vector pET-15b. Expression and purification of the protein was performed as described for Rab8a above, except that the polyhistidine tag was not removed for the pulldown experiment.

Removal of the His_6_-tag (Rab8a) was performed by overnight incubation at 4 °C with thrombin (GE Healthcare), followed by a second Ni^2+^-agarose column. The ‘flow-through’ fractions were collected, while the uncut proteins remained on the resin. Soluble aggregates were eliminated by running the sample through a Superdex 75 (16/60) gel filtration column (GE Healthcare) equilibrated in column buffer (20 mM Tris-HCl, 100 mM NaCl, 5 mM MgCl_2_, 1 mM DTT, pH 7.5). A peptide corresponding to residues 129-165 of RILPL2 was synthesized with an N-terminal hexahistidine tag (His_6_-RILPL2, Genscript). The peptide was solubilized in matching buffer with Rab (20 mM Tris-HCl, 100 mM NaCl, 5 mM MgCl_2_, 1 mM DTT, pH 7.5) prior to crystallization trials and calorimetry.

### Rab8a nucleotide exchange

For the pulldown with full-length RILPL2, nucleotide exchange was performed using purified WT Rab8a incubated in 10 mM EDTA for 10 minutes at room temperature in the presence of 10X molar excess GDP. The exchange was terminated by addition of 15 mM MgCl_2_ and excess nucleotides were removed by running samples through a PD10 column (GE healthcare), or by immediate gel filtration chromatography. To verify successful exchange, 100 μL the protein (>1mg/mL) was boiled for 10 min at 95 °C to denature the protein and release the nucleotide, followed by centrifugation for 30 min 16,000 x *g*, 4 °C to remove precipitated protein. The supernatant was mixed with running buffer (100 mM potassium phosphate, 8 mM thiobarbituric acid, pH 6.5) at a 1:1 ratio. The samples were loaded on an Acquity Ultra Performance system (Waters Corporation, Milford, MA, USA; or Varian 920 LC machine, Agilent, Stockport, UK) equipped with a ZORBAX 300SB-C18 column (Agilent, Stockport, UK). Elution profiles of GMP, GDP, GTP (Sigma Aldrich) and GppNHp (Jena Bioscience, Germany) were subjected to HPLC and compared with Rab8a. The nucleotide state of Rab8a(Q67L) was confirmed to be GTP-bound using the analytical HPLC strategy.

### *In vitro* kinase assays

For comparison of Rab8a phosphorylation by LRRK2 and MST3, kinase assays were performed with shaking at 30 °C for 3 h with concentrations as indicated of 970-end length LRRK2 WT or G2019S (PV4873 and PV4882 respectively, ThermoFisher) or GST-MST3 (supplied by MRC Reagents and Services, DU30889) and 2 µM of Rab8a 1-181 Q67L or Q67L+T72E as a negative control. The kinase reaction buffer is 50 mM Tris-HCl pH 7.5, 10 mM MgCl_2_, 150 mM NaCl, 2 mM ATP. Efficiency of Rab8a phosphorylation was compared using PhosTag gel electrophoresis and immunoblotting with Rab8a-pT72 antibody (Suppl Fig S1).

### Phosphorylation of Rab8a

It has recently been shown that the MST3 kinase can specifically and efficiently phosphorylate Rab8a at Thr72 *in vitro* (Vieweg, S et al manuscript in preparation). As MST3 is much easier to express than LRRK2, we decided to phosphorylate Rab8a at T72 using recombinant MST3. Full length GST-MST3 produced in insect cells (DU30889) was obtained from MRC-PPU Reagents and Services (https://mrcppureagents.dundee.ac.uk/reagents-proteins/overview). Rab8a was incubated with GST-MST3 at molar ratios between 4:1 to 9:1 (substrate:enzyme). Typical concentrations of Rab8a were 1-3 mg/ml, while the concentration of MST3 was 1 mg/ml in a total volume between 2-15 ml. The buffer of the reaction was adjusted to 50 mM Tris-HCl, 150 mM NaCl, 10 mM MgCl_2_ and 2 mM ATP, pH 7.5. The reaction mixture was incubated at room temperature overnight (12-18 hours). To separate pRab8a from the non-phosphorylated form, the reaction mixture was dialyzed against low-salt ion exchange buffer (10 mM MES, 10 mM NaCl, 5 mM MgCl_2_, 1 mM DTT, pH 5.2) for two hours and then loaded onto a MonoS 5/50 GL column (GE Healthcare) equilibrated to the low-salt ion-exchange buffer. Elution of pRab8a was performed by running a 50% gradient from low- to high-salt buffer (10 mM MES, 1 M NaCl, 5 mM MgCl2, 1 mM DTT, pH 5.2) over 30 column volumes. The phosphorylation of Rab8a_1-181_ was confirmed by PhosTag gel electrophoresis. In order to stabilize pRab8a, the pH was adjusted to pH 7.5 immediately after elution from the ion-exchange column.

### Crystallization, data collection and refinement

Crystals of pRab8(Q67L): His_6_-RL2 complex were obtained in a 1:1 molar ratio of protein:peptide at a total of 12 mg/mL. Crystals were grown in 100 mM HEPES buffer (pH 7), 10% PEG4,000, and 10% 2-propanol. Plate-like crystals were harvested in precipitant supplemented with 25% glycerol and stored frozen in liquid nitrogen. X-ray data were collected under a cryogenic nitrogen stream at 100K (beamline 24-ID-C, Advanced Photon Source).

Native diffraction data were reduced using XDS and aimless, followed by structure determination using the Phaser software in the PHENIX package [29, 30]. Initial rounds of molecular replacement using Rab8a (GppNHp, PDB code 4lhw) resulted in a solution for 2 molecules in the asymmetric unit. Following successful identification of Rab8a in the crystal lattice, the electron density for the coiled coil of the effector was apparent. Side chains for RILPL2 were clear in the initial electron density, and refinement was performed using multiple rounds of model building and energy minimization using PHENIX and COOT[31]. The asymmetric unit contains two molecules of Rab8a (A:4-176, B:2-176) bound to GTP and a magnesium ion, and two molecules of the effector (C:129-159, E:129-160). The hexahistidine tag at the N-termini of the effector is not seen in electron density maps, except for one histidine at the N-terminus of chain C.

### Structural analyses and superpositions

In general, structures were aligned using the ‘secondary structure matching’ (SSM) protocol in COOT. The backbone superpositions of Rab8a from multiple structures (complexed, uncomplexed) typically aligned with an RMSD of 0.4 Å. The heterotetrameric structures of Rab8a:RILPL2 and Rab7:RILP were aligned using the Superpose software in CCP4[32, 33]. In order to better visualize the relative positions of the effectors, secondary structures from all 4 molecules in each complex were aligned. A total of 330 residues were matched, including 21 residues from each chain of RILP and RILPL2. The overall RMSD for the backbone atoms was approximately 3 Å.

### Pulldown assays and isothermal titration calorimetry

Calorimetry was performed in triplicate on an ITC-200 instrument (GE Healthcare). Protein concentrations were calculated based on their Abs_280_ using a ND-1000 NanoDrop spectrophotometer (Thermo Scientific). Following purification of Rab8a, the protein was dialyzed together in the same buffer as RILPL2 (10 mM Tris-HCl, 300 mM NaCl, 5 mM MgCl_2_, 20 mM imidazole and 1 mM DTT). Samples were centrifuged at 13,200 rpm for 10 minutes prior to concentration determination and ITC analysis. The concentrations of proteins for injections were between 400-600 μM (His_6_-RILPL2, residues 129-165) and 40-60 μM Rab8a and pRab8a (1-181).

For *in vitro* pulldowns, full-length RILPL2 (1-211) was used. Rabs and RILPL2 (10 μM each) were mixed together in 1.5 mL centrifuge tubes with 25 µl Ni^2+^-agarose resin in a final volume of 1ml of binding buffer (20 mM Tris pH 8.0, 300 mM NaCl, 20 mM Imidazole, 5 mM MgCl_2_, 10 mM β-mercapotoethanol). The reaction mixture was subjected to mild shaking for 15 minutes. Following gentle centrifugation (1,000 rpm), the resin was washed 3 times with 1 ml of the binding buffer. Following release of proteins from resin with 50 µl elution buffer (20 mM Tris-Cl pH 8.0, 300 mM NaCl, 200 mM imidazole), samples were subjected to SDS-PAGE and visualization with 0.5% Coomassie Brilliant Blue.

### Plasmids for cellular assays

The plasmids used for co-immunoprecipitation experiments were acquired from MRC PPU Reagents and Services (https://mrcppureagents.dundee.ac.uk/reagents-proteins/overview): HA-empty pCVM5 (DU49303); GFP-empty pCDNA5 (DU13156); Flag-LRRK2 R1441G pCMV (DU13077); Flag-LRRK2 Y1699C pCMV (DU13165); HA-Rab8a WT pCMV (DU35414); HA-Rab10 WT pCMV (DU44250); RILPL2-GFP WT pCDNA5D FRT/TO (DU27481); RILPL2-GFP R130A pCDNA5D FRT/TO (DU68022); RILPL2-GFP R130Q pCDNA5D FRT/TO (DU27521); RILPL2-GFP R130E pCDNA5D FRT/TO (DU27520); RILPL2-GFP P131A pCDNA5D FRT/TO (DU68030); RILPL2-GFP P131C pCDNA5D FRT/TO (DU68031); RILPL2-GFP R132K pCDNA5D FRT/TO (DU68023); RILPL2-GFP R132A pCDNA5D FRT/TO (DU67110); RILPL2-GFP R132Q pCDNA5D FRT/TO (DU68037); RILPL2-GFP R132E pCDNA5D FRT/TO (DU27522); RILPL2-GFP F133A pCDNA5D FRT/TO (DU68033); RILPL2-GFP L135A pCDNA5D FRT/TO (DU68032); RILPL2-GFP R139A pCDNA5D FRT/TO (DU68025); RILPL2-GFP R139Q pCDNA5D FRT/TO (DU68024); RILPL2-GFP R139E pCDNA5D FRT/TO (DU68026); RILPL2-GFP K149A pCDNA5D FRT/TO (DU68029); RILPL2-GFP K149Q pCDNA5D FRT/TO (DU68027); RILPL2-GFP K149E pCDNA5D FRT/TO (DU68028); RILPL2-GFP E157A pCDNA5D FRT/TO (DU68036); RILPL2-GFP E157Q pCDNA5D FRT/TO (DU68034); RILPL2-GFP E157K pCDNA5D FRT/TO (DU68035), HA-Rab8a Q67L pCMV (DU39393), HA-Rab8a T22N (DU39392), JIP3-GFP pCDNA5D FRT/TO (DU27721), JIP4-GFP pCDNA5D FRT/TO (DU27684), RILPL1-GFP pCDNA5D FRT/TO (DU27305).

### Antibody reagents

Antibodies used in this study were diluted in 5% w/v bovine serum albumin in TBS supplemented with 0.1% Tween-20 and 0.03% w/v sodium azide. The Rabbit monoclonal antibody for total LRRK2 (N-terminus) was purified at the University of Dundee[34]. Anti-GFP (PABG1, Chromotek, used at 1:1000) anti-GFP (#2956, CST, used at 1:1000), anti-HA (3F10, Merck, used at 1:1000), anti-pT72-Rab8a (MJF-R20, Abcam, used at 0.5 µg/mL), anti-LRRK2 C-terminal (N241A/34, Neuromab, used at 1:1000), and anti-αTubulin (3873S, CST, used at 1:5000), anti-GAPDH (#sc-32233, Santa Cruz Biotechnology, used at 1:5000). Secondary antibodies used were Licor IRDye for 800CW goat anti-rabbit (925-32211), goat anti-mouse (926-32210) and 680LT goat anti-rat (925-68029) and goat anti-mouse (926-68020), all used at 1:10,000 dilution in TBS with 0.1% v/v Tween-20 (TBS-T) and horseradish peroxidase-conjugated rat IgG secondary antibody (#31470, Thermo Fisher Scientific) used at 1:10,000 dilution in 5% non-fat dry milk dissolved in TBS-T.

### Culture and transfection of cells

HEK293 cells were cultured in Dulbecco’s modified Eagle medium (Glutamax, Gibco) supplemented with 10% fetal bovine serum (FBS, Sigma), 100 U/ml penicillin and 100 µg/ml streptomycin. Transient transfections were performed 40-48 hr prior to cell lysis using polyethylenimine PEI (Polysciences) at around 60-70% confluence. Transfections for co-immunoprecipitation experiments were done in 10 cm round cell culture dishes using 3 µg of Flag-LRRK2 R1441G, 1 µg of HA control or HA-Rab8a and 1 µg of GFP control or GFP-RILPL2 cDNA construct per dish diluted in 1 mL of OPTIMEM media and supplemented with 20 µg of PEI and incubated for 20 min before being added to the cell media. 1 h before lysis cells were treated with 500 nM of MLI-2 inhibitor or 0.1% DMSO control. Lysates were clarified by centrifugation at 17,000 x *g* for 10 min.

### Co-Immunoprecipitation of Rab8a and RILPL2

Cells were washed with PBS and lysed in lysis buffer - 50 mM Tris-HCl pH 7.5, 1 mM EGTA, 10 mM sodium β-glycerophosphate, 50 mM sodium fluoride, 5 mM sodium pyrophosphate*10H_2_O, 0.27 M sucrose and supplemented fresh before lysis with 1% v/v Triton-x100, 1 tablet of cOmplete Mini (EDTA-free) protease inhibitor (Merck, 11836170001) per 10 mL of buffer, 0.1 µg/mL of microcystin and 1 µM of sodium orthovanadate.

For GFP immunoprecipitation, lysates were incubated with nanobody αGFP binder sepharose from MRC PPU Reagents and Services for 1 hr (15 µl of packed resin/0.5 mg of lysate). Bound complexes were recovered by washing the beads three times with wash buffer (50 mM Tris-HCl pH 7.5, 150 mM NaCl) before eluting with 2xSDS/PAGE sample buffer supplemented with 1% v/v 2-mercaptoethanol. The samples were denatured at 70 °C for 10 min and the resin was separated from the sample by centrifugation through a 0.22 μm Spinex column (CLS8161, Sigma).

### Gel electrophoresis and immunoblot analysis

Samples were run on gels consisting of a 4% w/v acrylamide stacking gel [4% w/v acrylamide, 0.125 M Tris-HCl pH 6.8, 0.2% v/v Tetramethylethylenediamine (TEMED) and 0.08% w/v ammonium persulphate (APS)] and 10% w/v acrylamide separating gel [10% w/v acrylamide, 0.375 M Bis-Tris pH 6.8, 1% v/v tetramethylethylenediamine (TEMED) and 0.05% w/v ammonium persulphate (APS)] in MOPS buffer (50 mM MOPS, 50 mM Tris, 1 mM EDTA, 0.1% w/v SDS) at 90-120 V. For Coomassie staining, gels were stained with InstantBlue™ Ultrafast Protein Stain (ISB1L, Sigma-Aldrich) according to the manufacturer’s instructions and the gels were imaged using LICOR Odyssey CLx. For immunoblot analysis, proteins were electrophoretically transferred onto nitrocellulose membranes (Amersham Protran 0.45 μm NC; GE Healthcare) at 90 V for 90 min on ice in transfer buffer [48 mM Tris/HCl, 39 mM glycine, 20% v/v methanol]. Transferred membranes were blocked with 5% w/v non-fat dry milk dissolved in TBS-T [20 mM Tris/HCl, pH 7.5, 150 mM NaCl and 0.1% v/v Tween 20] at room temperature for 1 h. Membranes were then incubated with primary antibodies overnight at 4 °C. After washing membranes in TBS-T 3×15 min, membranes were incubated with secondary antibodies at room temperature for 1 h. After washing membranes in TBS-T 3×15 min membranes were scanned using LICOR Odyssey CLx.

### PhosTag gel electrophoresis and immunoblot analysis

Samples were supplemented with 10 mM MnCl2 before loading gels. Gels for Phos-tag SDS/PAGE consisted of a stacking gel [4% w/v acrylamide, 0.125 M Tris/HCl, pH 6.8, 0.2% v/v tetramethylethylenediamine (TEMED) and 0.08% w/v ammonium persulfate APS] and a separating gel [10% w/v acrylamide, 375 mM Tris/HCl, pH 8.8, 75 μM PhosTag reagent (MRC PPU Reagents and Services), 150 μM MnCl_2_, 0.1% v/v TEMED and 0.05% w/v APS]. After centrifugation at 17,000 x g for 1 min, samples were loaded and electrophoresed at 90 V with the running buffer [25 mM Tris/HCl, 192 mM glycine and 0.1% w/v SDS]. For Coomassie staining, gels were stained with InstantBlue™ Ultrafast Protein Stain (ISB1L, Sigma-Aldrich) according to the manufacturer’s instructions and the gels were imaged using LICOR Odyssey CLx. For immunoblot analysis, gels were washed 3×10 min in 48 mM Tris/HCl, 39 mM glycine, 10 mM EDTA and 0.05% w/v SDS followed by one wash in 48 mM Tris/HCl, 39 mM glycine and 0.05% w/v SDS for 10 min. Proteins were electrophoretically transferred onto nitrocellulose membranes (Amersham Protran 0.45 μm NC; GE Healthcare) at 100 V for 180 min on ice in transfer buffer [48 mM Tris/HCl, 39 mM glycine, 20% v/v methanol]. Transferred membranes were blocked with 5% w/v non-fat dry milk dissolved in TBS-T [20 mM Tris/HCl, pH 7.5, 150 mM NaCl and 0.1% v/v Tween 20] at room temperature for 1 h. Membranes were then incubated with primary antibodies overnight at 4 °C. After washing membranes in TBS-T 3×15 min, membranes were incubated with horseradish peroxidase labelled secondary antibody diluted in 5% skimmed milk powder in TBS-T at room temperature for 1 h. After washing membranes in TBS-T (5×10 mins), protein bands were detected by exposing films (Amersham Hyperfilm ECL, GE Healthcare) to the membranes using an ECL solution (SuperSignal West Dura Extended Duration, Thermo Fisher Scientific).

## Data Availability

The co-ordinates for the structure of the pRab8a:RILPL2 complex have been deposited in the Protein Data Bank with accession code 6h3y. The co-ordinates will be released upon publication of the manuscript.

## Acknowledgments

We thank the excellent technical support of the MRC-Protein Phosphorylation and Ubiquitylation Unit (PPU) Cloning Service (Melanie Whightman), DNA Sequencing Service (coordinated by Gary Hunter), the tissue culture team (coordinated by Edwin Allen), Reagents and Services antibody and protein purification teams (coordinated by Hilary McLauchlan and James Hastie). This work was supported by Science Foundation Ireland Principal Investigator Awards (grant number 12/IA/1239 to ARK), Michael J. Fox Foundation for Parkinson’s research [grant number 6986 (to D.R.A.)]; the Medical Research Council [grant number MC_UU_12016/2 (to D.R.A.)]; the pharmaceutical companies supporting the Division of Signal Transduction Therapy Unit (Boehringer-Ingelheim, GlaxoSmithKline, and Merck KGaA, to D.R.A.). Data were collected at the Northeastern Collaborative Access Team beamlines, which are funded by the National Institute of General Medical Sciences from the National Institutes of Health (P41 GM103403). The Pilatus 6M detector on 24-ID-C beam line is funded by a NIH-ORIP HEI grant (S10 RR029205). This research used resources of the Advanced Photon Source, a U.S. Department of Energy (DOE) Office of Science User Facility operated for the DOE Office of Science by Argonne National Laboratory under Contract No. DE-AC02-06CH11357.

## Figure Legends

**Table 1.** Crystallographic Data and Refinement Statistics.

| **Table 1. Crystallographic Data and Refinement Statistics** |  |
| --- | --- |
| ***Data Collection*** |  |
| Beamline | NECAT APS, 24-ID-E |
| Wavelength (Å) | 0.9792 |
| Space group | P 2<sub>1</sub> 2<sub>1</sub> 2<sub>1</sub> |
| Cell [a, b, c, (Å)] | 60.333, 71.509, 114.784 |
| Resolution (Å) | 53.41 – 1.767 (1.83 – 1.767) |
| Total reflections | 324,590 (29,329) |
| Unique reflections | 49,201 (4,719) |
| Completeness (%) | 99.55 (96.33) |
| <I/σ> | 16.4 (1.69) |
| Multiplicity | 6.6 (6.2) |
| R<sub>merge</sub> | 0.06877 (0.9488) |
| R<sub>meas</sub> | 0.07469 (1.036) |
| R<sub>pim</sub> | 0.02883 (0.4097) |
| CC<sub>1/2</sub> | 0.999 (0.649) |
| ***Refinement*** |  |
| No. of reflections for R<sub>work</sub> | 49,193 (4,718) |
| No. of reflections for R<sub>free</sub> | 2,419 (241) |
| R<sub>work</sub> | 0.1789 (0.2882) |
| R<sub>free</sub> | 0.2105 (0.3049) |
| Number of non-hydrogen atoms | 3,832 |
| macromolecules | 3,401 |
| ligands | 72 |
| solvent | 359 |
| protein residues | 416 |
| RMSD bond lengths (Å) | 0.007 |
| RMSD bond angles (°) | 0.91 |
| Average overall B-factor | 36.54 |
| Mean B-factors (Å<sup>2</sup>)<br>protein/ligands/waters | 35.84/29.72/44.53 |
| Ramachandran analysis<br>favored/allowed (%) | 97.24/2.76 |
| PDB accession code | 6h3y |

Values in parentheses in the column on right correspond to the statistics in the highest resolution bin. RMSD, root-mean-square deviation.

**Expanded view Figure 1:**
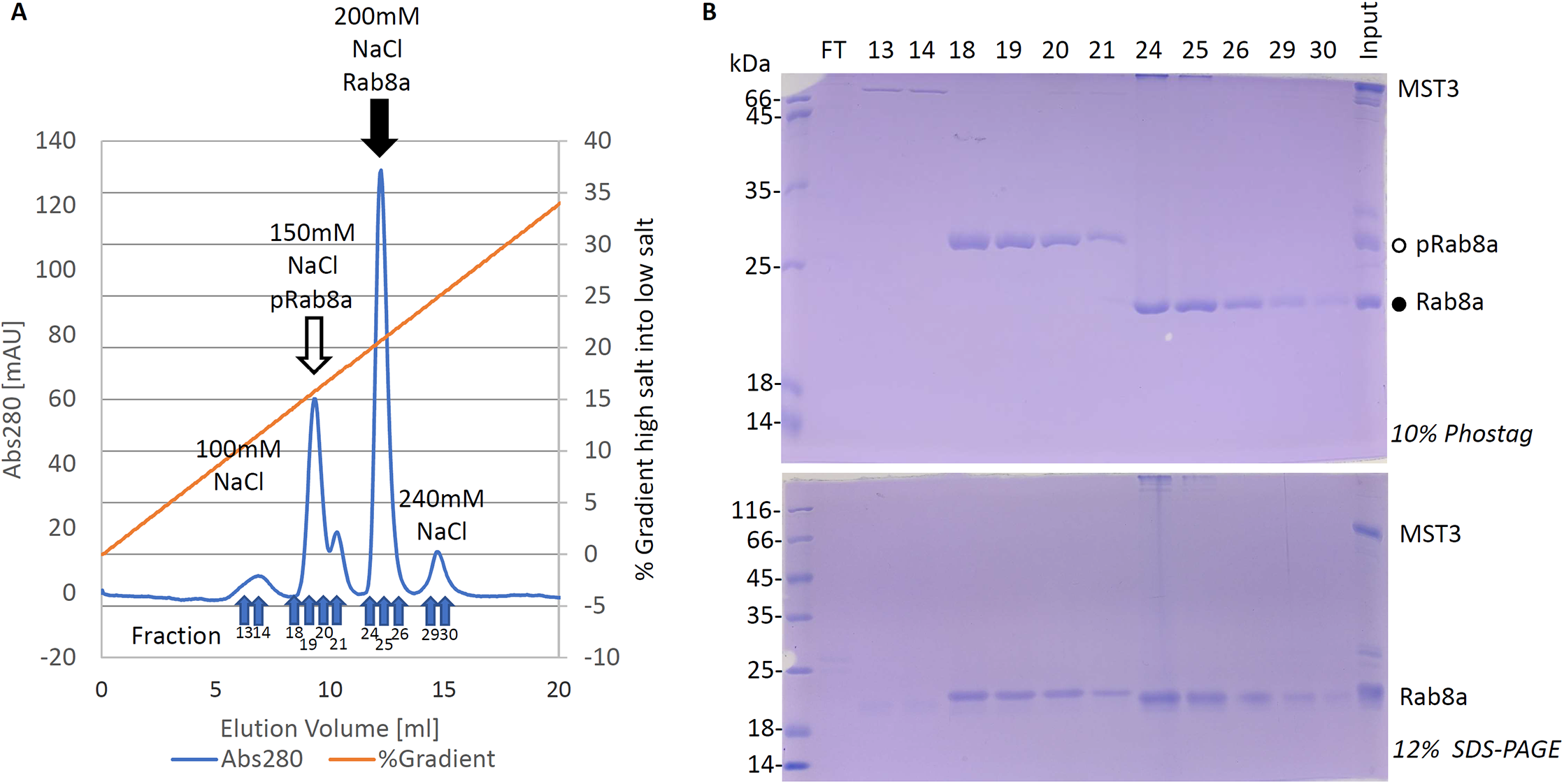
Purification of phosphorylated Rab8a. Following incubation of several mg of Rab8a_1-181_Q67L in the presence of GST-MST3 and ATP, phosphorylated Rab8a (pRab8a) was separated from the unphosphorylated protein by ion-exchange chromatography. **A**, Following dialysis in low salt buffer, reactants were loaded onto Mono-S cation exchange column (GE Healthcare) and eluted with a 50% gradient from low salt (10 mM) to high salt (1 M) buffered solution. During the salt gradient, 0.5 ml fractions were collected, and the two main peaks (arrows) corresponded to pRab8a and Rab8a. **B**, PhosTag gel (upper panel) and SDS-PAGE (lower panel) of the collected fractions. FT: unbound fraction *flow through*; Input: endpoint of phosphorylation reaction before loading the column. The PhosTag gel shows that the peak coming off at 150 mM NaCl contains the phosphorylated Rab8a.

**Expanded view Figure 3**: Figure legend same as for Figure 3

**Expanded view Figure 4:** Same figure legend as Fig 4B

**Expanded view Figure 5: JIP3 and JIP4 bind weakly to both Rab8a and LRRK2-phosphorylated Rab8a in cells.** Same figure legend as Fig 5D

**Supplementary Figure S1: Phos-tag analysis of MST3 mediated Rab8a phosphorylation.** Side by side analysis of wild type and G2019S insect cell expressed LRRK2 (residue 970-end) and full length insect cell expressed GST-MST3 mediated phosphorylation of recombinant Rab8a[Q67L, residues 1 to 181] or Rab8a[Q67L+T72E, residues 1 to 181]. Reactions were undertaken in the presence of 10 mM MgCl2 and 2 mM ATP for 3 hours at 37°C. Reactions were terminated by the addition of SDS sample buffer. Upper panel an aliquot of the reaction was analyzed by Phos-tag gel electrophoresis and protein visualized Coomassie Blue-staining. Bands corresponding to phosphorylated and non-phosphorylated Rab8a were marked with open (○) and closed (●) circles respectively. Lower panels samples were subjected to conventional gel electrophoresis and either stained with Coomassie (second panel) or subjected to immunoblot analysis using the LI-COR Odyssey CLx Western Blot imaging system with the indicated antibodies at 0.5-1 µg/mL concentration (two bottom panels). Similar results were obtained in at least two separate experiments.

